# Single-Molecule Localization Microscopy of 3D Orientation and Anisotropic Wobble using a Polarized Vortex Point Spread Function

**DOI:** 10.1101/2021.09.13.460135

**Authors:** Tianben Ding, Matthew D. Lew

## Abstract

Within condensed matter, single fluorophores are sensitive probes of their chemical environments, but it is difficult to use their limited photon budget to image precisely their positions, 3D orientations, and rotational diffusion simultaneously. We demonstrate the polarized vortex point spread function (PSF) for measuring these parameters, including characterizing the anisotropy of a molecule’s wobble, simultaneously from a single image. Even when imaging dim emitters (∼500 photons detected), the polarized vortex PSF is able to obtain 12 nm localization precision, 4-8° orientation precision, and 26° wobble precision. We use the vortex PSF to measure the emission anisotropy of fluorescent beads, the wobble dynamics of Nile red (NR) within supported lipid bilayers, and the distinct orientation signatures of NR in contact with amyloid-beta fibrils, oligomers, and tangles. The unparalleled sensitivity of the vortex PSF transforms single-molecule microscopes into nanoscale orientation imaging spectrometers, where the orientations and wobbles of individual probes reveal structures and organization of soft matter that are nearly impossible to perceive using molecular positions alone.

## Introduction

In the three decades since they were first detected optically in condensed matter, ^1–3^ single molecules (SMs) have become an invaluable tool for nanoscale imaging and spectroscopy across the biological, chemical, and material sciences. In particular, SM orientations can be used to quantify a variety of nanoscale biological and chemical processes, ^4^ such as reaction kinetics within porous catalytic nanoparticles; ^5^ nanoscale polymer deformation; ^6^ and the organization and higher-order structures of DNA, ^7,8^ amyloid fibrils, ^9,10^ lipid nanodomains, ^11^ and actin filaments. ^12,13^ By leveraging modern cameras, microscopes routinely collect highly sensitive, multiplexed data from each SM, i.e., using tens of pixels to estimate SM positions with nanoscale precision. Point-spread function (PSF) engineering, i.e., specifically designing an SM image to contain additional information beyond 2D position, has taken advantage of these multiplexed measurements for imaging SMs in 3D^14^ and for imaging molecular orientation in SM orientation-localization microscopy (SMOLM). ^4,10,11^ The field continues to evolve with theoretical studies to determine fundamental performance limits, ^15–17^ the assistance of machine learning to design optimal PSFs for 3D SMLM, ^18–20^ and new PSFs for measuring the 3D positions and 3D orientations of SMs. ^21–23^

The central challenge facing SM spectroscopists is the finite photon budget afforded to each individual fluorophore before irreversible photobleaching; due to photon shot noise, it is difficult to measure experimentally the many properties of SM emission, e.g., the position, arrival time, wavelength, and polarization of each incoming photon, simultaneously with high precision. Tradeoffs are unavoidable, and one’s sensitivity to a particular parameter can be improved to incredible degrees by reducing sensitivity to others, e.g., achieving ∼1 nm localization precision in MINFLUX by making the measurement largely independent of emission wavelength and polarization. ^24^ Here, we focus our attention on several outstanding questions in SMOLM. First, many existing methods have relatively poor sensitivity to molecular orientation along the polar direction *θ*, because their images of z-oriented molecules are relatively dim and diffuse compared to in-plane molecules. ^10,22^ Can we design a method with dramatically improved sensitivity to polar orientation? Second, existing techniques are largely limited to estimating rotational diffusion, or the “wobble” of an SM, as a scalar parameter averaged over all directions. ^16,17,21,23,25–27^ However, we know molecules wobble in full 3D space and, in certain environments, perhaps anisotropically, i.e., more in one direction versus another. ^28,29^ Is it possible to robustly quantify the degree of wobble anisotropy of an SM experimentally? Finally, what can we learn about the structure or organization of a biological sample through imaging the 3D orientations and wobble anisotropies of SMs with nanoscale resolution?

In this Article, we demonstrate a method, termed the polarized vortex PSF, for measuring the position, 3D orientation, and 3D rotational diffusion of single molecules. Here, we adopt the vortex phase mask, typically used for generating a donut-shaped illumination ring in STED^30^ and MINFLUX,^24^ and place it in the detection path of an epifluorescence microscope to create the PSF. We improve the orientation sensitivity of an existing implementation^23^ by splitting the detected fluorescence into two orthogonally polarized channels. The polarized vortex PSF dramatically changes shape with molecular orientation, thus enabling precise estimation of SM rotational diffusion in full 3D space using a single camera snapshot. We used the emission collected from fluorescent beads to verify the accuracy of our orientation and wobble measurements. We next measured the rotational dynamics of Nile red (NR) within lipid bilayers to quantify the condensing effects of cholesterol.^11^ Interestingly, the wobble anisotropies of NR change in response to its chemical environment, yielding a new method to elucidate intermolecular forces at work within the membrane. Finally, we used transient amyloid binding (TAB^31^) to image the orientations and rotational dynamics of Nile red in contact with amyloid fibrils,^10^ oligomers, and fibrillar tangles. We observe a diverse collection of NR orientation signatures that are correlated with aggregate morphology. To our knowledge, the polarized vortex PSF enables the first experimental demonstration of extracting seven parameters of information from the image of a single molecule: its 2D position (*x, y*), 3D orientation (*θ, ϕ* on a sphere), its wobble (*α, β*) in 3D, and its brightness *s*. While rotational diffusion tensors of molecular ensembles have been measured via nuclear magnetic resonance,^28^ we believe this is the first experimental report that quantifies the anisotropic rotational diffusion of a single molecule.

## Methods

### Modeling the rotational diffusion of a dipole-like emitter

We model a fluorescent molecule as a dipole-like emitter^32–34^ and represent its orientation with a unit vector ***µ*** = [*µ*_*x*_, *µ*_*y*_, *µ*_*z*_] ^T^ = [sin (*θ*) cos (*ϕ*), sin (*θ*) sin (*ϕ*), cos (*θ*)]^T^ (Figure 1A inset), where the superscript T denotes a matrix/vector transpose. Formally, ***µ*** describes the orientation of a molecule’s emission transition dipole moment at the instant it emits a fluorescence photon. Raw SMLM images contain typically hundreds to thousands of fluorescence photons from each SM, where each photon *h* may arise from a distinct orientation ***µ***_*h*_. Therefore, any SMOLM image contains information on a molecule’s mean orientation (*θ, ϕ*) and its rotational diffusion during the camera frame. This rotational diffusion can be assumed to be uniform within a hard-edged symmetric cone,^25,29^ or one can directly model rotational diffusion within an orientational potential well.^26,35^ If an SM’s rotational diffusion possesses sufficient symmetry, one may conveniently consider it as a mixture of two ideal sources: one immobilized dipole at a particular orientation plus one freely rotating molecule (i.e., an isotropic point source), each of a particular brightness.^7,26,27^

**Figure 1:**
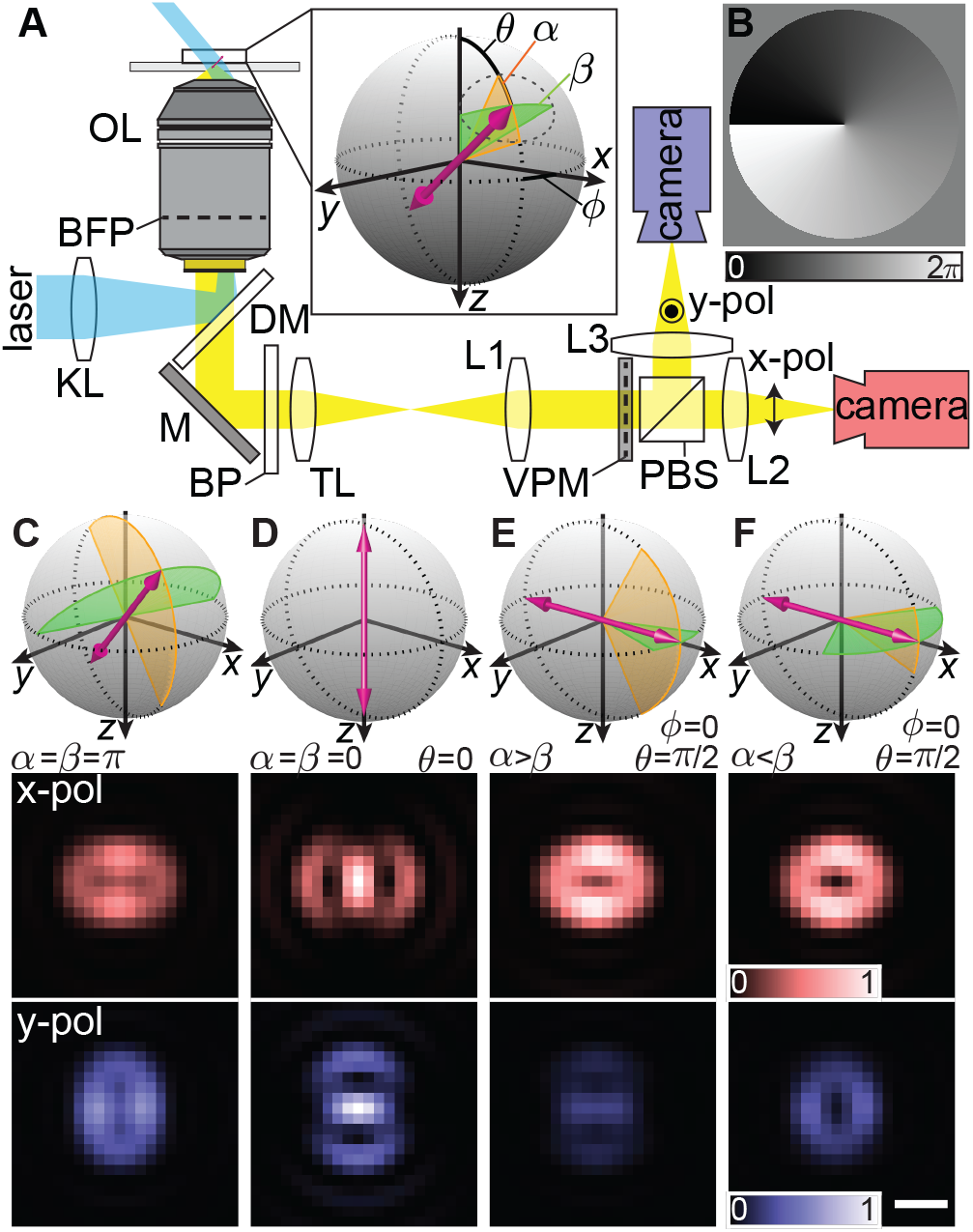
Single-molecule orientation localization microscopy (SMOLM) using the polarized vortex point spread function (PSF). A) Experimental setup. A circularly-polarized excitation laser (cyan) is coupled into an objective lens (OL) for highly-inclined illumination. Fluorescence (yellow) is collected by the same objective and isolated by a dichroic mirror (DM) and bandpass filter (BP) followed by a 4f optical system (lenses L1-L3, *f* = 150 mm) containing a vortex phase mask (VPM) and a polarizing beamsplitter (PBS). The VPM is placed in a conjugate back focal plane (BFP) to implement the vortex PSF. Two orthogonally polarized fluorescence images are captured by separate cameras or different portions of a single camera. Inset: The orientation of a transition dipole is parameterized by an azimuthal angle *ϕ* ∈ [− *π, π*), a polar angle *θ* ∈ [0, *π/*2], and an elliptical cone to model anisotropic rotational diffusion. This cone has aperture angles *α, β* ∈ [0, *π*], where *α* represents the extent of the cone’s aperture along the polar direction (orange), whereas *β* represents rotational wobble in the orthogonal direction (green). KL, widefield lens; M, mirror; TL, tube lens. B) Vortex phase mask. Color bar: phase (rad). C-F) Representative dipole orientations (top) and corresponding polarized vortex PSFs (bottom) for molecules with C) unconstrained and D) completely immobile rotational diffusion as well as anisotropic wobbles E) *α* = *π/*3, *β* = *π/*9 and F) *α* = *π/*9, *β* = *π/*3. Scale bar: 300 nm.

Here, we adopt a generalized model for rotational diffusion introduced by Backer and Moerner.^29^ During a camera frame, we assume a molecule’s orientation to be confined within a hard-edged elliptical region of the unit hemisphere, parameterized by two angles *α, β* ∈ [0, *π*] (Figure 1A inset). These two angles describe the extent of orientations that a molecule can explore along the polar direction (*α*) and the direction orthogonal to it (*β*). If *α* = *β*, then we term the rotational diffusion or wobble to be isotropic; else, we term it anisotropic. Note that we use the terms “anisotropic wobble” and “anisotropic rotational diffusion”^28^ intentionally, as our meaning is distinct from traditional fluorescence polarization anisotropy^36,37^ or rotational anisotropy. Polarization anisotropy refers to collecting photons in two or more polarized detection channels and is defined in terms of the fluorescence intensities collected from those channels. Polarization anisotropy has been combined with polarization modulation of excitation light to measure SM wobble.^8,38^ Here, wobble anisotropy directly quantifies the rotational dynamics of the molecule itself: the degree to which it is diffusing differently in two orthogonal directions.

To quantify the ellipticity of SM wobbling, i.e. rotational diffusion, we define the wobble anisotropy ratio *r*_*αβ*_ as

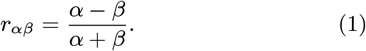

Since *α, β* ∈ [0, *π*], the ratio *r*_*αβ*_ varies from −1 to 1, similar to linear dichroism,^4,39,40^ and yields a straightforward interpretation of relative anisotropy. If |*r*_*αβ*_ |= 1*/*5, then *α* or *β* is 50% (1.5 times) larger than the other. Similarly, *α* or *β* is twice larger than the other if |*r*_*αβ*_ |= 1*/*3.

Assuming that a molecule’s rotational correlation time is faster than its fluorescence lifetime and the temporal resolution of an imaging sensor,^25,26,35^ its orientation trajectory can be fully characterized by a second-moment vector^41^ 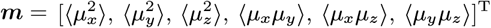, where each entry is a second moment of ***µ*** averaged over the camera integration time. Naturally, these orientational second moments contain information on both the 3D orientation of a molecule and its rotational diffusion. Given the first-moment orientation parameters (*θ, ϕ, α, β*) representing SM orientations within a hard-edged elliptical cone, one may calculate a 3 × 3 matrix ***M*** containing the orientational second moments, given by^29^

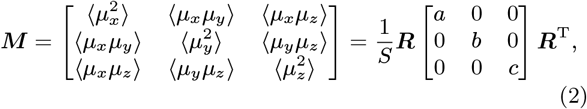

where

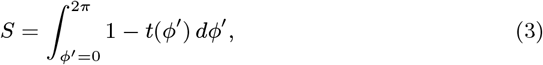

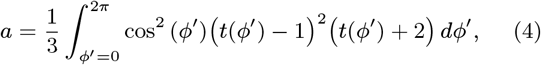

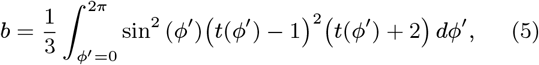

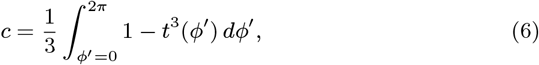

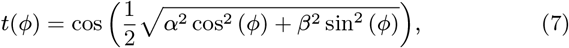

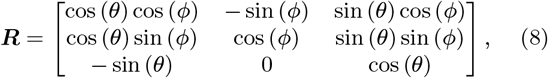

and *S* is the solid angle of the elliptical cone parameterized by *α* and *β* on the unit sphere (Figure 1A inset). The transformation matrix ***R*** rotates an elliptical cone aligned with the optical (*z*) axis to an arbitrary orientation (*θ, ϕ*), keeping the *α* axis of the ellipse oriented along the polar direction.

### Image formation and the polarized vortex PSF

An *n*-pixel image ***I*** ∈ ℝ^*n×*1^ of a dipole-like emitter can be modeled as a linear superposition of the six basis images ***B***_*d*_ of a fluorescence microscope weighted by the second-moment vector ***m***^17,29,41^ of the emitter, i.e.,

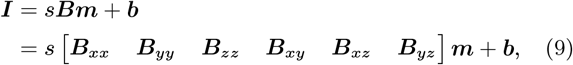

where *s* is the number of photons detected by the camera and ***b*** ∈ ℝ^*n×*1^ is the number of background photons in each pixel. The so-called basis images ***B***_*d*_ ∈ ℝ^*n×*1^, where *d* ∈ *{xx, yy, zz, xy, xz, yz}*, can be calculated using vectorial diffraction theory.^29,41–43^ Importantly, these basis images are determined only by the design of the imaging system.

As a consequence of the linearity of the imaging model, the basis images ***B***_*d*_ directly affect the performance of the optical system for measuring each orientational second-moment *m*_*d*_.^17^ Since the orientation (*θ, ϕ*) and rotational diffusion (*α, β*) parameters of an SM are directly related with the second moments ***m*** (Eqns. 2-8), we desire each ***B***_*d*_ to be unique and have approximately equal energy to one another. If these images do not have these properties, then certain SM orientations could become indistinguishable and/or measurement precision could suffer greatly.^22^

To achieve these goals, we introduce the polarized vortex PSF, which can be implemented by inserting a vortex phase mask (VPM, Figure 1B) and a polarizing beamsplitter (PBS) into a 4f optical system attached to a standard epifluorescence microscope (Figure 1A). We used a liquidcrystal spatial light modulator to create the vortex phase mask within a bespoke 4f system with two polarized imaging channels, as described previously.^31,44^ An unpolarized vortex PSF was recently reported by Hulleman and coworkers^23^ for simultaneously imaging the 3D positions and orientations of SMs. While this method is easy to implement because it does not require two polarized imaging channels, it shows reduced sensitivity for measuring the orientational second moments in terms of the Cramér-Rao bound (Figure S3). Thus, for dim SMs, resolving anisotropic wobble is difficult without polarization-sensitive imaging (Figure S2).

The polarized vortex PSF clearly resolves freely rotating molecules (Figure 1C) from fixed molecules (Figure 1D), as well as an SM wobbling largely in the *α* direction (Figure 1E) vs. one wobbling mostly in the *β* direction (Figure 1F). In fact, the polarized vortex PSF provides superior precision for measuring the second moments ***m*** compared to several state-of-the-art orientation-sensing PSFs, including the polarized standard PSF,^10^ bifocal microscope,^45^ CHIDO,^21^ and unpolarized vortex PSF.^23^ This performance holds when molecules are near a refractive index interface as well as when they are within a matched medium (Figure S3).

### Preparation and imaging of fluorescent beads

Fluorescent beads (Thermo Fisher Scientific, FluoSpheres, 0.1 µm, 580/605, F8801) were used in this work for calibrating a home-built microscope (Supporting Notes 1, 2) and characterizing the measurement performance of the polarized vortex PSF. The beads were spin-coated on an ozone-cleaned coverslip (Marienfeld, No. 1.5H, 22 × 22 mm, 170 ± 5 µm thickness) and excited by a 561-nm laser source. In the experimental demonstration, we collected 100 images of each bead with a 20 ms camera exposure time and observed approximately 30,000 photons per bead (Table S1). The bead positions and orientations were estimated using a maximum-likelihood estimator (Supporting Note 3) as described below.

### Preparation and imaging of supported lipid bilayers

Supported lipid bilayers (SLBs) were prepared as described previously.^11^ Briefly, 25 mg/mL dipalmitoylphosphatidylcholine (di(16:0) PC, DPPC, Avanti Polar Lipids, 850355) with and without 40% mol/mol cholesterol (Sigma-Aldrich, C8667) in chloroform were first dried overnight in vacuum. The lipids were then re-suspended in Tris buffer (100 mM NaCl, 3 mM Ca^2+^, 10 mM Tris, pH 7.4) to a 1 mM lipid concentration. A 25-passage extrusion (Avanti Polar Lipids, 610000) was performed on a hot plate (95°C) after 30-min vortex mixing under nitrogen. The monodispersed DPPC large unilamellar vesicles (LUVs) were next placed onto ozone-cleaned coverslips and incubated in a water bath at 65°C for 1 hour to form SLBs. After 30 min cooling to 21°C, the coverlips were rinsed with Tris buffer (100 mM NaCl, 10 mM Tris, pH 7.4) thoroughly for SMOLM imaging. Nile red (NR, Thermo Fisher Scientific, AC415711000) in Tris buffer was added onto the coverslips and excited by the 561-nm laser (5.42 kW ·cm^−2^ intensity at the sample) in our system. Image stacks of 4,000 frames with 50 ms exposure were recorded in these experiments. NR photon statistics observed from PAINT-SLB imaging are shown in Figure S13 and Table S1.

### Preparation and imaging of amyloid aggregates

The 42 amino-acid residue amyloid-*β* peptide (A*β*42) was synthesized and purified by Dr. James I. Elliott (ERI Amyloid Laboratory, Oxford, CT) and prepared as described previously.^10^ In short, the purified amyloid powder was re-lyophilized after suspension in hexafluoro-2-propanol (HFIP) and sonication at room temperature for one hour. To further purify the monomeric amyloid precursors, the lyophilized A*β*42 was dissolved in 10 mM NaOH, sonicated for 25 min in a cold water bath and filtered first through a 0.22 µm and then through a 30 kD centrifugal membrane filter (Millipore Sigma, UFC30GV and UFC5030).

To prepare amyloid aggregates, we incubated 10 µM monomeric A*β*42 in phosphate-buffered saline (PBS, 150 mM NaCl, 50 mM Na_3_PO_4_, pH 7.4) at 37 °C with 200 rpm agitation for 43-70 hours (fibrils and oligomers) and 13 days (fibril networks with globular tangles). The aggregated structures were adsorbed to ozone-cleaned cell culture chambers (Cellvis, C8-1.5H-N, No. 1.5H, 170 ± 5 µm thickness) for 1 hour followed by a PBS rinse for removing residual aggregates. A PBS solution (200 µL) containing 50 nM Nile Red (Fisher Scientific, AC415711000) was added for TAB SMOLM imaging. Image stacks of 10,000 frames with 20 ms camera exposure under the same 561-nm laser excitation (0.88 kW ·cm^−2^ intensity at the sample) were recorded in these experiments. NR localization and orientation statistics are shown in Table S1 and S2.

### SMOLM image analysis

Before analyzing experimental data, we calibrated our image analysis algorithms by imaging fluorescent beads and performing phase retrieval (see Supporting Note 2 and Figure S12). Background fluorescence within each camera frame was first estimated using spatial and temporal filtering (see Supporting Note 3). Next, the brightness *s*, 2D position (*x, y*), and second moments ***m*** of individual SMs within each imaging frame were jointly estimated using a bespoke sparsity-promoting maximum-likelihood estimator (RoSE-O^46,47^). This estimation was followed by a weighted least-square estimator for projecting the estimated second moments ***m*** to angular parameters (*α, β, θ*, and *ϕ*, Supporting Note 3). Typical analysis of a SMOLM dataset (131 × 262 pixels × 5,000 frames) took approximately 2 hours on a 32-core workstation (AMD Ryzen™ Threadripper™ 3970X CPU with 64 GB RAM running MATLAB R2021a).

## Results and Discussion

### Measurement Performance in Simulations and in Fluorescent Bead Imaging

We first validate the performance of the polarized vortex PSF using synthetic images of SMs with various orientations and rotational diffusion (See Supporting Note 4 for more details). Our Monte Carlo studies indicate that our statistical estimation algorithm (RoSE-O) efficiently detects SM images of the polarized vortex PSF at practical low signal-to-background ratios (SBRs, 510 photons detected and 2.3 background photons per pixel, Figure S4). Further, our algorithm achieves good precision and accuracy for measuring location and brightness (precision: *σ*_*x,y*_ = 12.5 nm, *σ*_*s*_ = 36 photons, accuracy: *x* − *x*_0_ = *y* − *y*_0_ = 0.0 nm, *s* − *s*_0_ = 2 photons), as well as orientation (precision: *σ*_*θ*_ = 0.025*π* rad = 4.5°, *σ*_*ϕ*_ = 0.043*π* rad = 7.7°, accuracy: *θ* − *θ*_0_ = 0.002*π* rad = 0.4°, *ϕ* − *ϕ*_0_ = 0.0 rad = 0.0°). Importantly, the PSF and algorithm are able to resolve orientations in full 3D, i.e., throughout the entire orientation sphere, without degeneracy (Figures S5-S8).

For rotational diffusion (*α, β*), our estimator exhibits a slight negative bias (*α* − *α*_0_ = −0.056*π* = −10.1°, *β* − *β*_0_ = −0.069*π* = −12.5°) when molecular diffusion is present *α*_0_ *>* 0, *β*_0_ *>* 0). This bias stems from molecules appearing to be fixed in orientation (*α* = 0 or *β* = 0) even though the true rotational diffusion is not zero (9,893 of 31,734 trials for SBR = 510 photons detected and 2.3 background photons per pixel, Figure S9A). Interestingly, when this error occurs, only one wobbling parameter is biased (*α* = 0 or *β* = 0 but not both), and other estimates are precise and accurate (precision: *σ*_*x,y*_ = 12.2 nm, *σ*_*s*_ = 36 photons, *σ*_*θ*_ = 0.025*π* rad = 4.5°, *σ*_*ϕ*_ = 0.041*π* rad = 7.4°, accuracy: *x* − *x*_0_ = *y* − *y*_0_ = 0.0 nm, *s* − *s*_0_ = 2 photons, *θ* − *θ*_0_ = 0.004*π* rad = 0.7°, *ϕ* − *ϕ*_0_ = 0.0 rad = 0.0°). In rare cases, our estimator measures a completely fixed molecule (*α* = *β* = 0) if the true rotational diffusion is nonzero (57 of 31,734 trials for the situation shown in Figure S9A).

Notably, our investigations reveal that these estimation inaccuracies are consistent with unavoidable biases stemming from photon shot noise.^27^ Errors can be reduced by increasing the SBR and eventually completely removed once the SBR reaches 5,000 photons detected and 1.0 background photons per pixel (Figure S9). Since it is extremely unlikely in practical scenarios that the rotational diffusion of an SM is completely zero in one direction and non-zero along another direction, we remove all localizations with *α* = 0 ∧ *β >* 0 or *α >* 0 ∧ *β* = 0 in this work. This filtering process improves both the accuracy and precision of measuring SM rotational diffusion at the cost of reducing the number of SM localizations (the remaining 79% of the localizations provides precision: *σ*_*α*_ = 0.145*π* rad = 26.1°, *σ*_*β*_ = 0.148*π* rad = 26.7°, *σ*_*rαβ*_ = 0.243, accuracy: *α* −*α*_0_ = −0.023*π* rad = −4.1°, *β* −*β*_0_ = −0.011*π* rad = −2.0°, *r*_*αβ*_ −*r*_*αβ*0_ = −0.018, Figures S10 and S11).

Next, we experimentally validate the imaging concept by measuring the emission pattern of fluorescent beads. These nanospheres are typically assumed to behave as ideal point sources, especially those that are smaller than the diffraction limit (∼*λ/*2NA ≈ 250 nm), and are widely used for calibrating a variety of single-molecule imaging systems.^48–50^ Figure 2A shows representative polarized vortex PSF images of 100-nm beads spin-coated on glass coverslips and embedded in immersion oil. Given our imaging model [Equation (9)] and the orientational second moments of an isotropic emitter 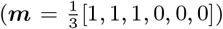, we expect these images to resemble an equal linear combination of the ***B***_*xx*_, ***B***_*yy*_, and ***B***_*zz*_ basis images (Figure 2B). Each bead image indeed contains a mixture of these features, including the double-lobed ring pattern of ***B***_*xx*_ in the *x* channel, a ring within ***B***_*yy*_ in the *y* channel, and a low-contrast “hole” at the center of each donut effected by the bright center spot of ***B***_*zz*_ in both channels.

**Figure 2:**
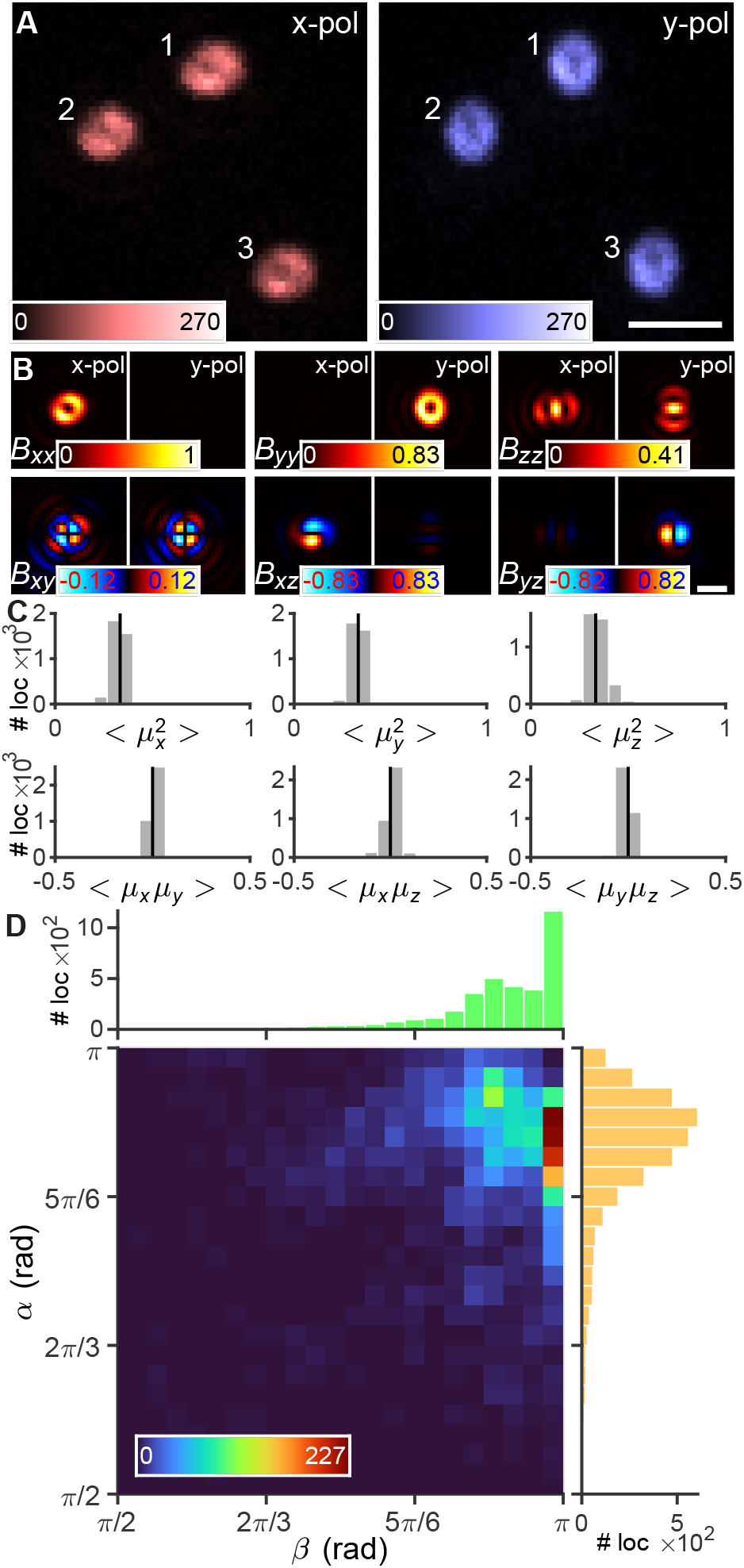
Emission anisotropy of fluorescent beads. A) Representative raw fluorescence images. Color bars: photons/pixel. B) Aberration calibrated orientational basis images. Images are normalized in each basis. Color bar: normalized intensity. C) Estimated orientational second moments of 3,752 localizations collected from 38 individual beads. Black lines indicate the second moments for an ideal isotropic emitter (*α* = *β* = *π*). D) Measured wobbling angles *α* and *β* for each bead localization shown in (C). The examples shown in (A) exhibit (Bead 1) *α* = 0.95*π, β* = 0.93*π*, (Bead 2) *α* = 0.95*π, β* = 0.92*π*, and (Bead 3) *α* = 0.90*π, β* = 0.98*π*. Color bar: localizations per histogram bin. Histograms depict distributions of (right) *α* and (top) *β*. Scale bars: (A) 1 *µ*m, (B) 500 nm.

Fitting images of each bead using RoSE-O yields second-moment distributions close to that of an ideal isotropic emitter (Figure 2C). While the fluorophores within these nanoparticles are likely fixed in position and orientation, their collective emission pattern is consistent with a single dipole undergoing large rotational diffusion (*α* = 0.88*π* ±0.083*π* rad, *β* = 0.93*π* ± 0.081*π* rad, mean ± std). Interestingly, the experimental *α* and *β* distributions show smaller mean values and larger standard deviations compared to Monte Carlo simulations (*α* = 0.93*π* ± 0.047*π* rad and *β* = 0.99*π* ± 0.027*π* rad for a ground-truth emitter with *α*_0_ = *β*_0_ = *π* and brightness matching the experiment, *s*_0_ = 30,300 ± 10,400 photons). Thus, we observe anisotropy and variations thereof that are consistent with non-uniform distributions of fluorophores within each sphere^51^ and between spheres, respectively; these non-uniformities are detectable due to the high 3D orientation sensitivity of the polarized vortex PSF.

### Resolving Anisotropic Rotational Diffusion of SMs within Supported Lipid Bilayers

Lipid membranes are essential cellular building blocks useful for organizing many processes.^52,53^ Their heterogeneous compositions and dynamic architectures, embodied by structures like cholesterol-enriched nanodomains, make them compelling targets for nanoscale imaging.^54^ We have previously utilized the Tri-spot^51^ and Duo-spot PSFs to resolve the composition of and enzyme activity within supported lipid bilayers (SLBs) using SMOLM.^11^ Compared to other engineered PSFs, the polarized vortex PSF has enhanced sensitivity to all six orientational second moments (***m***) and a relatively small footprint.^23^ Here, we used Nile red (NR) PAINT^55^ (points accumulation for imaging in nanoscale topography) and the polarized vortex PSF to estimate the orientations and rotational diffusion of single molecules within supported lipid bilayers (SLBs). In PAINT, single NR molecules spontaneously bind to and unbind from lipid membranes, emitting fluorescence flashes while they are within a non-polar environment.

We imaged SLBs composed of dipalmitoylphosphatidyl-choline (DPPC, di(16:0) PC) with (Figure 3A) and without cholesterol (chol) (Figure 3B). Representative vortex PSF images of NR show a dramatic change influenced by chol concentration (Figure 3C,D). Our orientation measurements show that NR exhibits a more out-of-plane orientation in DPPC without chol (*θ* = 0.31*π* ± 0.11*π* rad) than it does when embedded within DPPC with 40% chol (*θ* = 0.21*π* ±0.12*π* rad). In accordance with our previous observations, chol-induced condensation within the membrane orients NR molecules more parallel to the acyl chains of neighboring lipid molecules.^11^

**Figure 3:**
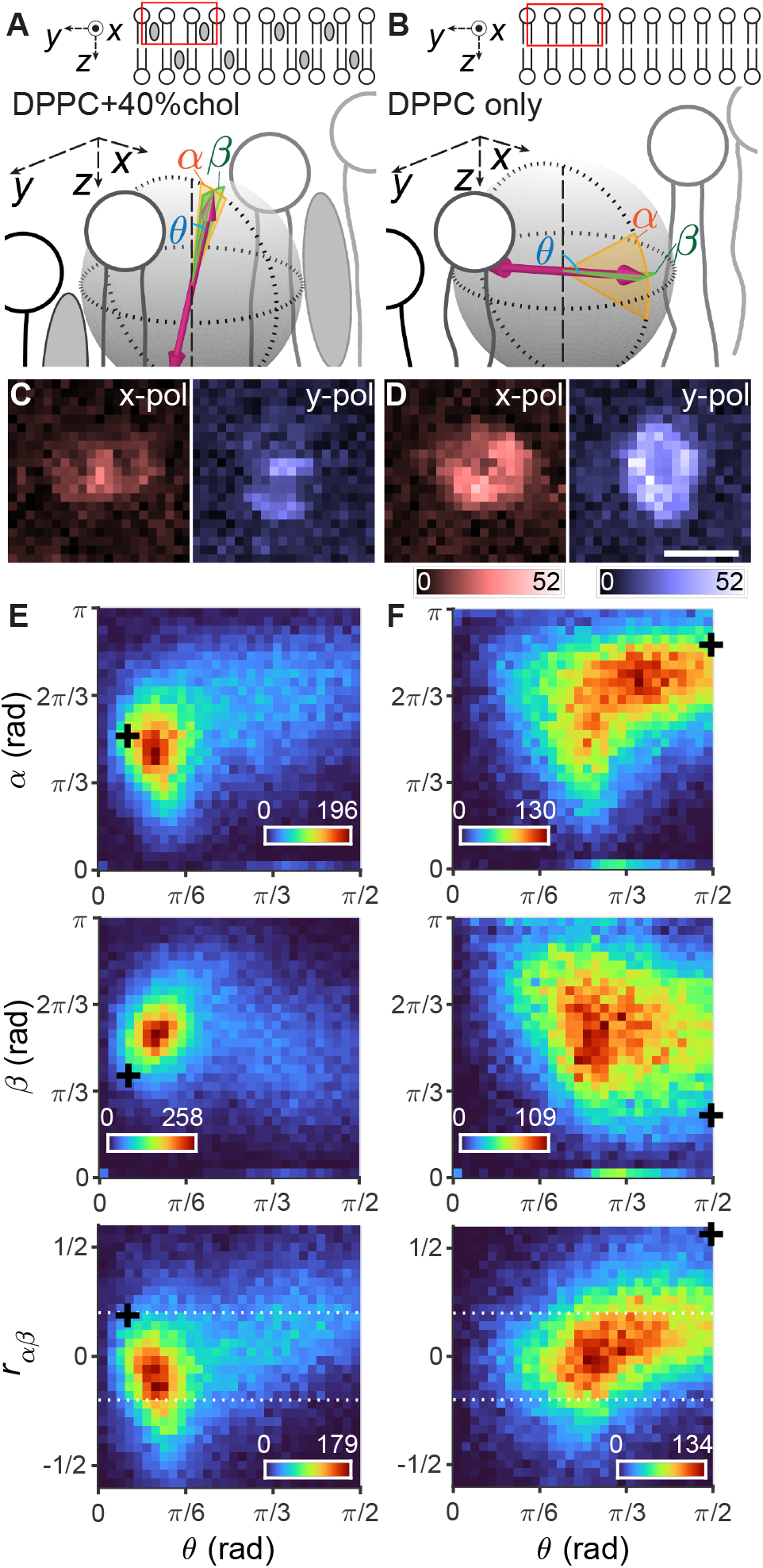
PAINT SMOLM of the 3D orientations and rotational diffusion of single Nile red (NR) molecules in supported lipid bilayers (SLBs). A,B) Schematic of the orientation and rotational diffusion of NR dipoles within DPPC bilayers A) containing 40% and B) no cholesterol (chol). Gray ovals represent chol in the SLBs. C,D) Representative images of single NR captured in x- and y-polarized channels within C) DPPC+40% and D) DPPC-only SLBs. Color bars: photons/pixel. E,F) Estimated NR rotational diffusion angles (top) *α* and (middle) *β* and (bottom) anisotropy ratio *r*_*αβ*_ vs. orientation *θ* in E) DPPC+40% and F) DPPC-only SLBs. Black crosses correspond to the estimated rotational diffusion and orientations of the examples shown in (B,C). White dashed lines denote |*r*_*αβ*_ |= 1*/*5 where *α* or *β* is 50% larger than the other. Color bars: localizations per histogram bin. Scale bar: 500 nm.

The presence of chol also affects NR’s rotational diffusion *α* along the polar direction and causes more confined wobbling when chol is present (*α* = 0.54*π* ± 0.19*π* rad, Figure 3E top) compared to the DPPC-only situation (*α* = 0.63*π* ± 0.20*π* rad, Figure 3F top). Note that the afore-mentioned ensemble-averaged mean values of *α* hide the considerable correlation between diffusion *α* and polar angle *θ* of NR at the SM level. Interestingly, chol-induced ordering has less impact on molecular wobble *β* perpendicular to the polar direction (*β* = 0.54*π* 0.18*π* rad for DPPC+40% chol vs. *β* = 0.57*π* ±0.22*π* rad for DPPC only). We observe that the distribution of wobble *β* and orientation *θ* is extremely broad for NR in DPPC without chol (Figure 3E,F middle).

Our measurements of anisotropy ratio *r*_*αβ*_ for NR within the two different SLBs are similar when averaged across the ensemble (DPPC+40% chol: *r*_*αβ*_ = 0.0 ± 0.27 vs. DPPC only: *r*_*αβ*_ = 0.058 ± 0.25). However, a key advantage of the polarized vortex PSF is its ability to quantitatively measure the anisotropy ratio and orientation simultaneously of each observed SM. When chol is present, the rotational wobble of NR is slightly greater in the *β* direction or nearly isotropic, i.e., equal in all directions (Figure 3E bottom). However, *r*_*αβ*_ increases substantially in the DPPC-only SLB, especially for highly tilted NR (*r*_*αβ*_ = 0.16 ± 0.23 for *θ* ≥7*/*18*π*, Figure 3F bottom).

The polarized vortex PSF’s ability to detect this anisotropy in molecular wobble yields remarkable insight into how NR interacts with its surroundings; NR is strongly affected by intermolecular forces exerted by the crowded forest of lipid tails, even when its residence time within the membrane is short (mean fluorescence burst length of 16 ms, Figure S13, Table S1). When it is aligned with the SLB surface, it readily collides with neighboring acyl chains and exhibits nearly 40% larger wobble in the polar direction compared to the orthogonal direction, corresponding to *r*_*αβ*_ = 0.16. This anisotropy completely changes character when NR is oriented perpendicular to the membrane, regardless of the presence of chol.

### Visualizing the Structural Organization of Amyloid Assemblies

A defining feature of amyloid diseases, such as Alzheimer’s and Parkinson’s disease, is the aggregation of proteins into fibrils and their deposition into plaques and intracellular inclusions. There are more than 50 different proteins or peptides, e.g., amyloid-*β* and *α*-synuclein, that are known to aggregate.^56,57^ Regardless of the specific amino acid sequence, the aggregation process begins with misfolded monomers that nucleate the growth of small soluble oligomeric intermediates into fibrillar assemblies.^58,59^ Although many amyloid aggregates share a common structural motif, e.g., the cross-*β* sheet,^60^ oligomers and fibrils are highly heterogeneous in both structure and size and not all aggregation intermediates are equally cytotoxic.^61,62^ In particular, the cross-*β* architecture of amyloid fibrils exhibit *β*-sheet strands that run perpendicular to the long axis of the fibril with grooves formed by its amino-acid side chains.^63^ These grooves serve as binding sites of amyloid-sensitive fluorescent molecules, such as NR and Thioflavin T.^64,65^

In addition to sensing hydrophobicity,^66^ we hypothesize that the rotational diffusion of NR molecules, while bound transiently to surfaces of amyloid aggregates, may be used to quantify the organization of the stacked *β* sheets. Here, we incubated recombinant amyloid-*β* (1-42) peptides (A*β*42) on ozone-cleaned coverslips to form amyloid fibrils (see Methods for further details). We observed fluorescence bursts from individual NR molecules localized to both fibrils and punctate oligomers (Figures 4A, S14A-C and Movie S1), a signature of transient amyloid binding (TAB).^31,67^ Standard SMLM reconstructions clearly resolve morphological differences between fiber-like and small sphere-like structures but are unable to discern further detail due to limited spatial resolution.

**Figure 4:**
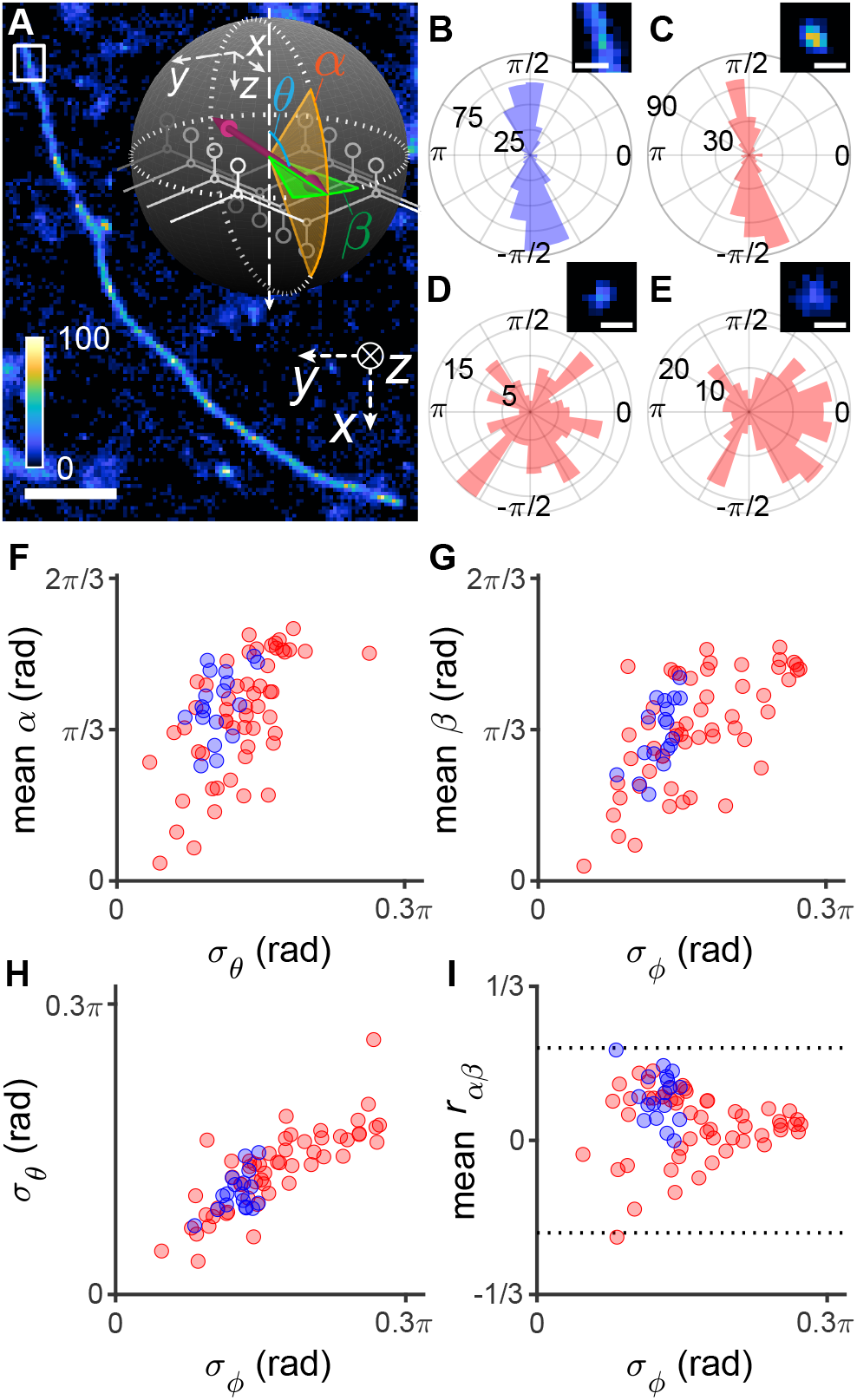
Characterizing NR binding orientations and rotational diffusion on amyloid-*β* (A*β*42) fibrils and oligomers using transient amyloid binding (TAB) SMOLM. A) Single-molecule localization microscopy (SMLM) image of A*β*42 aggregates. Inset: Schematic of the orientation and rotational diffusion of an NR dipole transiently bound to a cross-*β* assembly. Color bar: localizations per bin (40 × 40 nm^2^). B-E) Histograms of measured NR azimuthal orientations *ϕ* B) within the fibril region denoted by the white box in (A) and C-E) on representative small spherical amyloid aggregates. Insets: SMLM images corresponding to each histogram using the same color scale as (A). F-I) 3D orientations and rotational diffusion of NR on (blue) 18 thin fibril segments (∼200 nm in length) and (red) 54 small oligomeric structures. F) Mean polar wobble *α* vs. standard deviation of polar angle *θ*. G) Mean wobble angle *β* vs. standard deviation of azimuthal angle *ϕ*. H) Standard deviation of *θ* vs. standard deviation of *ϕ*. I) Mean anisotropy ratio *r*_*αβ*_ vs. standard deviation of *ϕ*. Black dotted lines denote |*r*_*αβ*_ |= 1*/*5. Scale bars: (A) 1 *µ*m, (B-E) 150 nm. Localization statistics are reported in Table S1

By carefully analyzing images from the polarized vortex microscope, we can further measure the 3D orientation and rotational diffusion of each NR molecule. As observed previously,^10,65^ azimuthal orientations *ϕ* show that individual NR molecules bind mostly parallel to the long axis of the fibrils (Figure 4B, std. dev. *σ*_*ϕ*,B_ = 0.14*π*). Further, NR molecules bound to the thin extended fibril shown in Figure 4B lie mostly parallel to the coverslip (*θ*_B_ = 0.40*π* ±0.08*π* rad) and exhibit slight anisotropic wobbling (*α*_B_ = 0.38*π* ± 0.30*π* rad, *β*_B_ = 0.29*π* ± 0.25*π* rad, *r*_*αβ*,B_ = 0.13 ± 0.25). While variations in wobble angles (*α, β*) are broad relative to their difference (*α* −*β*), we still observe significant wobble anisotropy for a majority of molecules on fibrillar structures, especially for those parallel to the coverslip (more than 65% of localizations with *θ* ≥ 7*π/*18 exhibit *r*_*αβ*_ *>* 0, Figure S14F). This anisotropy corresponds to more rotational freedom in the polar direction (Figure 4A inset), implying that the cross-*β* grooves significantly hinder NR from wobbling in the coverslip plane.

In contrast to thin fibrils, NR binds to small oligomers in various ways depending on the specific particle. While appearing oligomeric, some small aggregates show fibrillar character in terms of NR orientations, exhibiting well-aligned azimuthal orientations *ϕ* (Figure 4C, *σ*_*ϕ*,C_ = 0.16*π*, see Figure S14D for more examples) and nearly in-plane polar orientations (*θ*_C_ = 0.42*π* ± 0.08*π* rad). These aggregates also have fibril-like wobble anisotropy (Figure 4C: *α*_C_ = 0.38*π* ± 0.28*π* rad, *β*_C_ = 0.31*π* ± 0.26*π* rad, *r*_*αβ*,C_ = 0.11 ± 0.25). On the other hand, other small aggregates show more diverse azimuthal orientations (Figure 4D,E, *σ*_*ϕ*,D_ = 0.27*π, σ*_*ϕ*,E_ = 0.27*π*, see Figure S14E for more examples) and out-of-plane polar orientations (Figure 4D,E: *θ*_D_ = 0.36*π* ± 0.11*π* rad, *θ*_E_ = 0.35*π* ± 0.10*π* rad). Interestingly, NR exhibits mostly isotropic wobbling on this type of oligomer (Figure 4D,E: *α*_D_ = 0.50*π* ± 0.26*π* rad, *β*_D_ = 0.47*π* ± 0.26*π* rad, *r*_*αβ*,D_ = 0.03 ± 0.29, *α*_E_ = 0.51*π* ± 0.26*π* rad, *β*_E_ = 0.48*π* ± 0.28*π* rad, *r*_*αβ*,E_ = 0.04 ± 0.30).

We analyzed, in total, 54 small oligomeric structures (149 avg. localizations per structure) and compared their estimated NR orientations and rotational diffusion against those of 18 thin fibril segments (252 avg. localizations per segment). We first observed positive correlations between the wobble angles *α* and *β* of each localization and the standard deviations in 3D orientation (*σ*_*θ*_ and *σ*_*ϕ*_) across all nearby localizations (within a 100 nm radius), regardless of whether NR was bound to fibrils or oligomers (Figure 4F-H). That is, the rotational diffusion of each individual molecule tended to be large when the ensemble variations in SM orientation *θ* and *ϕ* were large. The orientations and rotational diffusion of NR on oligomeric structures exhibit a variety of distributions (Figure 4F-H red), while NR bound to fibril segments shows a specific orientational character (mean *σ*_*θ*,fibril_ = 0.105*π* ± 0.020*π*, mean *σ*_*ϕ*,fibril_ = 0.127*π* ± 0.017*π*, mean *α*_fibril_ = 0.39*π* ± 0.07*π*, mean *β*_fibril_ = 0.32*π* ± 0.07, mean *r*_*αβ*,fibril_ = 0.096 ± 0.053, Figure 4F-H blue).

Fibrillar structures with well-aligned *β*-sheet assemblies show obvious anisotropic rotational diffusion (*r*_*αβ*_ *>* 0, Figure 4I blue). On the other hand, NR molecules on oligomeric structures (Figure 4I red) experience both types of anisotropic rotational diffusion (*r*_*αβ*_ *<* 0 and *r*_*αβ*_ *>* 0) when NR orientations are well-aligned (small *σ*_*ϕ*_). This anisotropy gradually disappears as NR orientations become more varied (large *σ*_*ϕ*_), suggesting that *β*-sheet stacking is largely disordered within these small aggregates. We note that SMOLM data clearly distinguish between more ordered, “fibril-like” oligomeric structures^65^ (Figures 4C and S14D) vs. disordered, amorphous oligomers (Figures 4D,E and S14E), even for amyloid structures aggregated under identical conditions within the same field of view. Differences between these particles cannot be resolved in standard SMLM reconstructions (Figures 4C-E and S14D,E insets).

When accumulating NR wobble measurements across all fibrils (Figure S14F) and across all oligomers (Figure S14G), we find distinct NR motions compared to those within lipid membranes (Figure 3E,F). We observe a non-negligible amount of fixed NR (*α* = *β* = 0) in contact with both fibrils and oligomers (Figure S14F-G, left and middle). Interestingly, the mean anisotropy ratio *r*_*αβ*_ is largely the same for NR on fibrils (*r*_*αβ*_ = 0.054 ± 0.27, Figure S14F right) vs. oligomeric structures (*r*_*αβ*_ = 0.040 ± 0.27, Figure S14G right). These data show the critical necessity of using SM and single-particle techniques in amyloid studies; particle-to-particle variations and heterogeneity cannot be detected with ensemble averaging.

Since the polarized vortex PSF measures precisely the orientations and rotational diffusion of individual probe molecules, we next explore how SMOLM may be used to resolve and characterize the complex organization of bundled and intersecting fibrils within a large amyloid network. After a longer aggregation time (13 days), SMLM shows an example aggregate network consisting of isolated single fibrils, thicker bundles of fibrils, and globular tangles (Figure 4A). Strikingly, we observed that the rotational anisotropy *r*_*αβ*_ of NR varies dramatically throughout the network. NR molecules bound to fibrillar tangles exhibit large anisotropy (*r*_*αβ*_ ≥1*/*5), especially when the estimated polar angles are large (NR parallel to the coverslip, magenta arrows in Figure 5B). On the other hand, for NR inclined out of the coverslip plane (small *θ*), junctions between globular tangles and fibril bundles show more isotropic wobbling (*r*_*αβ*_ ≈0, white arrows in Figure 5C). These data are evidence that the polarized vortex PSF is able to resolve the full 3D orientations of *β*-sheet grooves (Movie S2).

**Figure 5:**
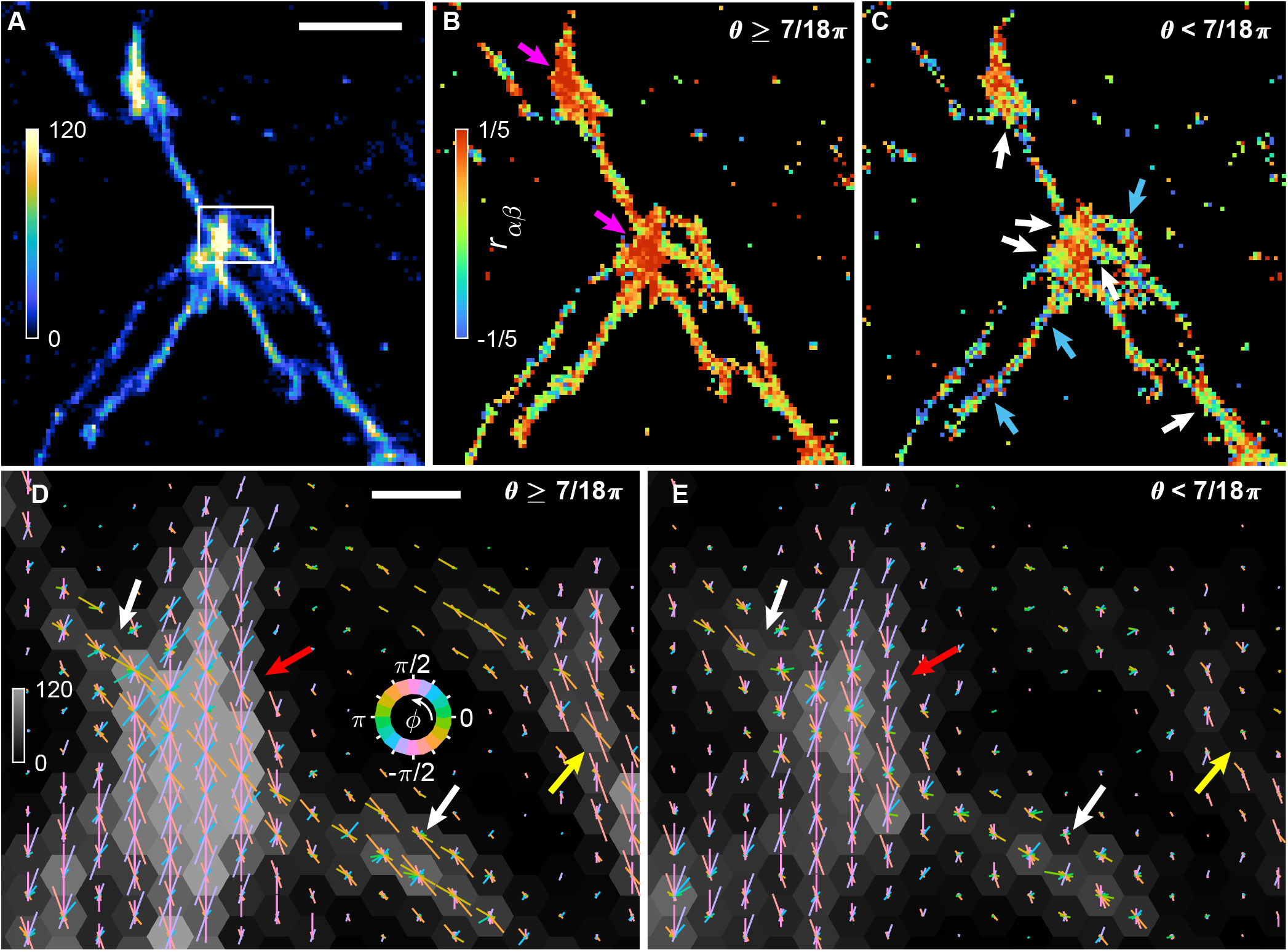
Mapping NR binding orientations and rotational diffusion throughout an amyloid fibrillar network using TAB SMOLM. A) SMLM image. Color bar: localizations per bin (40 × 40 nm^2^). B,C) SMOLM images color-coded by mean anisotropy ratio *r*_*αβ*_ for NR molecules oriented B) parallel to the coverslip (*θ* ≥ 7*π/*18 rad = 70°) and C) out-of-plane (*θ <* 7*π/*18 rad = 70°). Magenta and blue arrows denote regions with large wobble *α* in (B) and large wobble *β* in (C), respectively, while white arrows denote isotropic wobble in (C). Color bar: mean anisotropy ratio *r*_*αβ*_ per bin (40 × 40 nm^2^). D,E) 2D hexagonal histogram (grayscale) of NR localizations oriented D) in-plane (*θ* ≥ 7*π/*18 rad) or E) out-of-plane (*θ <* 7*π/*18 rad) within the white box in (A). Distributions of NR azimuthal orientations *ϕ* within each spatial bin (46.2 nm center-center distance) are superimposed as polar histograms (bin size Δ*ϕ* = *π/*9 rad). Longer line segments represent greater numbers of localizations oriented along the *ϕ* direction of each line. Yellow arrows label a well-ordered fibril while white arrows denote a relatively disordered fibril. Red arrows depict an amyloid tangle. Gray color bar: localizations per bin. Multicolor circular color bar: azimuthal orientation *ϕ*. Scale bar: (A) 1 *µ*m, (D) 100 nm.

TAB SMOLM also detected a significant population of NR with small polar angles and isotropic or anisotropic rotational diffusion (blue arrows in Figure 5C); notably, some molecules exhibited strong *β*-angle wobbling (negative *r*_*αβ*_). While the SMLM reconstruction (Figure 5A) suggests that these regions are fibrils, both the NR orientations and wobble anisotropies here differ dramatically from our observations of single thin fibers adhered to coverglass (Figure 4F). Instead, the NR orientation data imply that these are fibrillar bundles with crossed-*β* grooves lying both parallel and inclined relative to the coverslip. While these bundles would be extremely difficult to detect by 3D SMLM without exquisite (∼few nm) spatial resolution, TAB SMOLM is able to directly resolve them through the orientational binding behaviors of NR.

One particular advantage of TAB SMOLM over traditional bulk polarization anisotropy is its ability to map fluorophore orientations with SM sensitivity. Therefore, we may analyze the full population distribution of orientation data within each region of the amyloid network to better understand its architecture. We have isolated one such region in the white box in Figure 5A. In Figure 5D,E, we show polar histogram maps of azimuthal orientation *ϕ*, accumulating localizations within each hexagonal spatial bin. Longer line segments represent greater numbers of localizations oriented along the *ϕ* direction of each line.

We first note that the large tangle (aforementioned magenta arrow in Figure 5B) contains *β*-sheet grooves oriented along many different azimuthal directions, both for grooves parallel to the coverslip (red arrow in Figure 5D) and inclined away from the coverslip (red arrow in Figure 5E). These data agree with our intuition about the overall structure of these tangles; they contain a complex, interconnected network of fibers extending in all directions in 3D.

These orientation maps also resolve differences in the architectures of fibrillar bundles. Two bundles near the tangle contain NR localizations whose azimuthal orientations lie parallel to the long axis of the bundles (white and yellow arrows in Figure 5D), as expected. Both bundles also contain some crossed-*β* grooves oriented out of the plane. One bundle’s grooves are mostly aligned azimuthally with the long axis of the bundle, indicating a well-ordered bundle extending along the coverslip (yellow arrow in Figure 5E). However, the bundle intersecting the tangle contains grooves oriented in many azimuthal directions (white arrows in Figure 5E). We hypothesize that these orientations may be a signature of early-stage tangling within the fibrillar bundle. The differences in the organization of these fibrillar bundles are not resolvable by standard SMLM.

## Conclusions

We report the polarized vortex PSF as a powerful tool for precisely and accurately estimating the 2D positions, 3D orientations, and 3D rotational “wobble” of single fluorophores. The PSF is more compact than most other engineered PSFs, thereby improving its ability to detect weak emitters and enabling denser SM imaging. In addition, its high sensitivity to all six orientational second moments allows it to readily discriminate between rotational motion along the optical axis versus that in the perpendicular direction. These capabilities permit experimenters to map single-molecule rotational dynamics across a wide field of view with nanoscale resolution and exquisite measurement precision.

In this study, the polarized vortex PSF resolved SM dynamics in two biological model systems. First, NR within DPPC SLBs shows a wide variability in rotational wobble but tends to exhibit more rotational diffusion along the direction parallel to acyl chains (*α*) compared to the orthogonal direction (*β*). In contrast, chol-induced ordering of the lipid molecules causes NR wobbling to become more isotropic and less varied between NR binding events. Second, we found that NR has distinct orientations and rotational diffusion characteristics depending on whether it is in contact with extended fibrils or sphere-like oligomers. In particular, fibrils are characterized by 1) well-ordered NR azimuthal orientations *ϕ* (i.e., small standard deviations *σ*_*ϕ*_ over an ensemble of localizations), 2) an intermediate amount of rotational diffusion (*α, β*) at the SM level, and 3) a modest degree of NR anisotropic wobble *r*_*αβ*_. In contrast, between various oligomers, we observe large variations in SM wobble anisotropy *r*_*αβ*_ that mirror a wide variety of NR azimuthal orientation distributions, ranging from ordered (i.e., fibril-like) to disordered (large *σ*_*ϕ*_). These orientation data provide insights into *β*-sheet organization that cannot be resolved by conventional SMLM.

We note that given their 3-10 nm thickness,^68^ it is extremely difficult to resolve the architecture of fibrils and fibrillar bundles using most 3D SMLM techniques. However, recent innovations in interferometric SMLM,^69^ measuring fluorescence lifetimes of SMs near a metal or graphene interface to resolve axial positions,^70,71^ and 3D MINFLUX^72^ could achieve the necessary spatial resolution. Here, we adopt an orthogonal approach, namely measuring SM orientational dynamics, because these dynamics reveal how biomolecules are organized in a way that complements traditional localization microscopy. The photon-efficient sensitivity of the polarized vortex PSF to SM orientation (*θ, ϕ*) and anisotropic wobble (*α, β*) resolves the organization of amyloid fibrils and tangles with just 500-700 photons detected on average per molecule.

To realize the full potential of SMOLM, optical PSF designs must continue to improve detection and measurement sensitivity, while also extending operable imaging depths and mitigating challenges from optical aberrations and auto-fluorescence. In addition, more robust and expedient image analysis tools, including machine learning, for measuring SM position, orientation, and wobble simultaneously will facilitate broader adoption of SMOLM technology. In the future, we believe SMOLM will join fluorescence emission and excited state lifetime spectroscopies as a useful tool for mapping the organization of molecules in the condensed phase, as well as for measuring intermolecular forces within crowded environments. We look forward to the fourth decade of single-molecule spectroscopy with excitement and anticipation.

## Acknowledgement

This work is supported by the National Science Foundation under grant no. ECCS-1653777 and by the National Institute of General Medical Sciences under grant no. R35GM124858 to M.D.L. The authors thank Jin Lu for preparation of the supported lipid bilayers.

## Supporting Information Available

The following files are available free of charge.

- Supporting Information: Additional experimental notes on the optical instrumentation, procedure of optical-aberration retrieval, analysis and processing of SMOLM images, and generation of synthetic data. Supporting figures providing characterization of the polarized vortex PSF and comparison of the Cramér-Rao bound of different PSFs; Monte Carlo simulations evaluating the performance of the polarized vortex PSF with the RoSE-O estimator; retrieved optical aberrations in the polarized imaging system; quantitative characterization of NR blinking in SLBs and additional orientation measurements of NR molecules on amyloid oligomers and fibrils. Table comparing localization, orientation, and rotational diffusion statistics of SMOLM data estimated by the polarized vortex PSF.
- Supporting Movies: Transient amyloid binding (TAB) SMOLM using the polarized vortex PSF and SMOLM reconstruction of Nile red 2D positions and 3D orientations throughout an amyloid fibrillar network.
- Supporting Data: Supplementary data is available via OSF (https://osf.io/fu7gd/?view_only=6b16cadd11174a57adf44f4ca0ecc1e1) and by request.

## Graphical TOC Entry

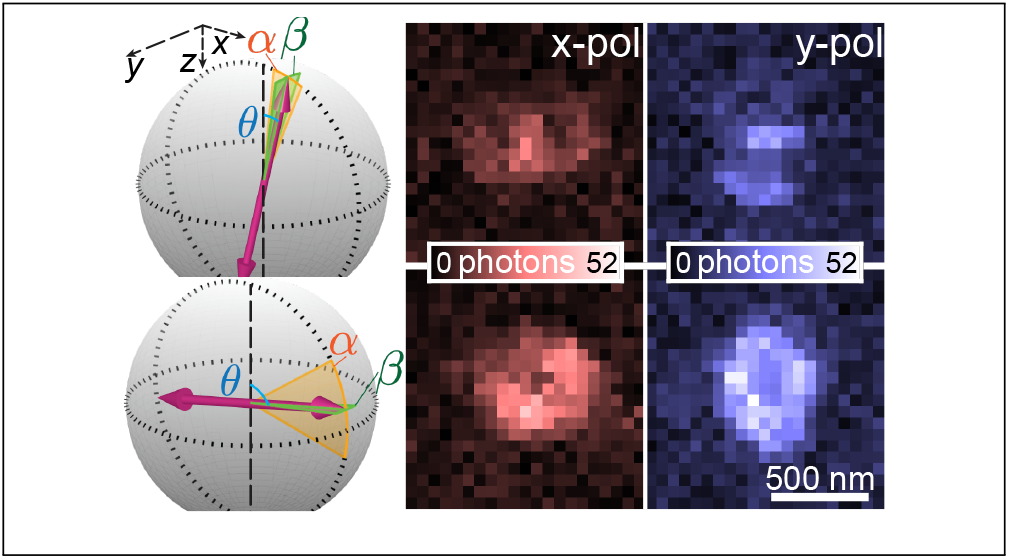

